# Predicting neutron radiation exposure characteristics from an *in vitro* human skin model using RNA-seq

**DOI:** 10.64898/2026.01.29.702673

**Authors:** Myles W. Gardner, Danielle S. LeSassier, Joshua C. Gil, Jesse C. Werth, Madeleine N. Pont, Guy Garty, Elizabeth A. Scheuermann, Helen C. Turner, Patrick J. Cocola, Charles Grice, Vivi-ana M. June, Catie A. Vaccaro, Brooke E. Tashner, F. Curtis Hewitt

## Abstract

Accurate assessment of low-dose neutron radiation exposure remains a central challenge in biodosimetry, particularly for applications requiring non-invasive sample types such as skin. Here, we characterized the transcriptional response of a three-dimensional *in vitro* human skin model (EpiDermFT) to neutron irradiation at doses up to 0.75 Gy, measured from pre-exposure through a 14-day post-exposure period. RNA sequencing revealed greater than 800 significantly altered genes, including upregulation of *FOS, FOSB, CDKN1A, MDM2*, and *GADD45A*, and downregulation of *NRG1, H3C11*, and *CENPX*. Gene ontology enrichment indicated activation of DNA damage checkpoint signaling, cell cycle arrest, and stress-response pathways, alongside suppression of nucleosome assembly and DNA replication processes. Machine learning models trained on transcriptomic features exhibited strong predictive performance across biodosimetric endpoints. Classification models accurately distinguished irradiated from sham samples (AUC > 0.99), and regression models achieved high accuracy for estimating both absorbed dose (R^2^ = 0.97) and days post-exposure (R^2^ = 0.99). The latter, while highly predictive, may partially reflect transcriptional shifts associated with progressive degradation of the *in vitro* tissue model over time. Collectively, these findings demonstrate that RNA-based molecular signatures from human skin tissue provide a robust framework for quantitative estimation of neutron radiation exposure and temporal response dynamics.

## Introduction

Exposure to low dose ionizing radiation (LDIR) can occur through multiple routes. This includes occupational exposures, including the nuclear, medical, and air travel industries, more generally through medical procedures (e.g., computed tomography scans), natural sources (e.g., cosmic rays), or during a nuclear incident, either accidental or intentional. Given the prolonged exposure individuals may have and studies highlighting the risk of LDIR exposure,^1–7^ understanding the biological impact and signatures of neutron radiation is critical for both assessing long term impacts on health and overall exposure status.

Biodosimetry provides a way to measure and predict radiation exposure through biological changes. Biomarker identification for biodosimetry often focuses on signatures from invasive collection matrices, such as blood or plasma.^8–10^ Gold standard methods for estimating exposure, like the dicentric chromosome assay (DCA), are time consuming and require substantial expertise to review.^11,12^ Collecting invasive samples presents challenges in a mass exposure scenario and the results may not sufficiently capture indicators of LDIR. Skin is the first line of exposure and presents an alternative to more traditional sample types that can be easily collected through non-invasive means, such as swabbing. Understanding the molecular changes occurring in the skin and the ability of associated biomarkers to support predictive modeling may provide tools to help determine low dose exposure and associated health risks.

RNA is a commonly targeted biomarker for biodosimetry efforts, given the rapid expression changes that can occur following exposure. Studies focused on impacts of LDIR have observed increased expression for pathways associated with survival and protection, such as double strand break repair, antioxidant defense, and proteolysis, over pathways more commonly associated with apoptosis and cycle progression that is typically seen at higher doses.^13–16^ However, these studies primarily looked at gene expression within minutes to hours post-exposure.^14,15,17^ While understanding the immediate impacts of radiation is important, this timing is less realistic in real world scenarios, either in a triage event or part of normal screening. The short-term focus makes it difficult to understand the longer-term impact of LDIR on gene regulation and expression trends. Additionally, many of these studies use x-ray as the ionizing radiation source and the translatability to neutron exposure at low doses is unclear.

LDIR studies are made challenging by the limited models available and studying the biological response in skin is no different. Mouse models are limited by the strain-based variability of radiation responsiveness, known differences in cellular response compared to humans, and structural difference in the skin architecture,^18–20^ making translation of potential markers to humans challenging. Human two-dimensional (2D) cell cultures are another standard model for assessing radiation impact but lack the complexity of tissue, reducing the number of useful biomarkers that can be observed, and do not fully capture the interplay of the stratified matrix.^21,22^ Three-dimensional (3D) human skin equivalent models present an *in vitro* alternative that better captures the heterogeneity and architectural composition of human skin. EpiDerm Full Thickness (FT) (MatTek Corporation) is one such 3D model composed of differentiated human-derived normal epidermal keratinocytes (NHEK) and normal dermal fibroblasts (NHFB) forming a multi-layered *in vitro* tissue. This model has been used for a range of dermal focused studies from toxicity assessments, wound healing, carcinogenesis, and non-neutron radiation exposure making it a useful and reproducible model.^15,23,24^

In the current study, we used the 3D human skin model, EpiDermFT, to identify neutron LDIR-responsive expression signatures over a two-week post-exposure timeframe using RNA sequencing (RNA-seq). This study extends the timeframes for investigating LDIR responses beyond the typical 24-hour assessment. Samples were exposed to various low doses of neutron radiation, ranging from 0.00 to 0.75 Gy. The resulting signatures were leveraged to build predictive models for exposure, dose, and the days post-exposure to better assess the value in biodosimetry for neutron LDIR.

## Results

### EpiDermFT RNA Extraction

To determine the optimal RNA extraction conditions a small batch shipment of 12 EpiDermFT tissues were maintained and collected. Multiple conditions were tested, including variable post-collection storage, lysis conditions, and extraction kits. Details and results of extraction development can be found in Supplemental Materials and Results. Extraction methods were assessed against RNA-seq requirements, including a minimum of 500 ng total RNA yield and a RIN value of greater than or equal to 7.0 for RNA quality. Ultimately, preservation in RNA*later* paired with physical homogenization in RLT buffer containing β-mercaptoethanol followed by extraction with the Qiagen RNeasy Plus Miniprep kit gave the best results (2,097.0 ± 127.3 ng RNA; RIN 7.2 ± 0.1). This extraction method was then used for all neutron study samples. Table 1 shows the average RNA yield by condition using this extraction workflow. Individual samples were then prepared and submitted for RNA-seq.

**Table 1:**
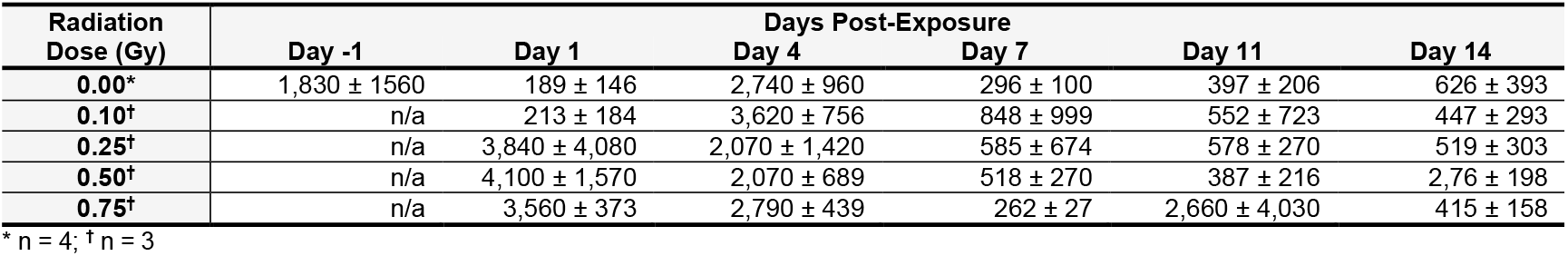
Average RNA yields (ng) from extracted EpiDermFT samples.

### RNA-sequencing Results

After quality control read trimming, 8.96 × 10^9^ reads remained with an average 5.34 ± 1.20 × 10^7^ reads per sample. The average Phred score was 35.44 ± 0.14. The reads were successfully mapped to the human reference genome and 21,533 unique ensemble gene IDs were assigned.

### Differential Expression and Signature Discovery Results

Differential expression between the exposed samples and sham controls yielded 832 total genes that had significant regulation changes (adjusted p-value < 0.05), of which 320 had an absolute log_2_ fold-change greater than 0.25 (137 up-regulated and 183 down-regulated). The differential expression, when broken down by cumulative dose and day post-exposure show unique trends (Table 2). The number of differentially expressed genes increased substantially once the dose increased to 0.50 Gy. Regarding day post-exposure, there is large increase in the number of differentially expressed genes from 62 on day 1 to 255 on day 4. There was a notable decrease in the number of differentially expressed genes after 14 days when only 16 genes showed any significant changes in regulation.

**Table 2:** Number of significantly (adjusted p-value < 0.05) up- or down-regulated genes with an absolute log_2_fold change greater than 0.25, summarized by dose and days post-exposure.

| Radiation Dose (Gy) | Gene Count | Days Post-Exposure | Gene Count |
| --- | --- | --- | --- |
| 0.10 | 49 | 1 | 62 |
| 0.25 | 33 | 4 | 255 |
| 0.50 | 100 | 11 | 205 |
| 0.75 | 93 | 14 | 16 |

The differential expression between sham and exposed samples for each of these genes is illustrated in the volcano plot in Figure 1A. A total of 832 genes showed significant differential expression (adjusted p-value < 0.05) between sham and exposed samples, and among these, *FOS* and *FOSB* had the largest fold-change magnitude. For complete comparison of sham vs. exposed samples see Table S1. Both *FOS* and *FOSB* were selected for modeling exposure and dose, as they showed clear, positive correlations between cumulative dose and normalized abundance at all time points (Figure 1B). Similar dose-dependent expression changes were also observed in many other genes identified during feature selection. However, the kinetics greatly varied across genes. For example, *CDKN1A* was up-regulated at higher doses of neutron radiation, but the effects on this gene were less prominent more than one-week post-irradiation. *NRG1*, on the other hand, was consistently down-regulated in all irradiated samples. For a complete break-down of dose- and day-dependent differentially abundant genes see supplemental Table S2.

**Figure 1:**
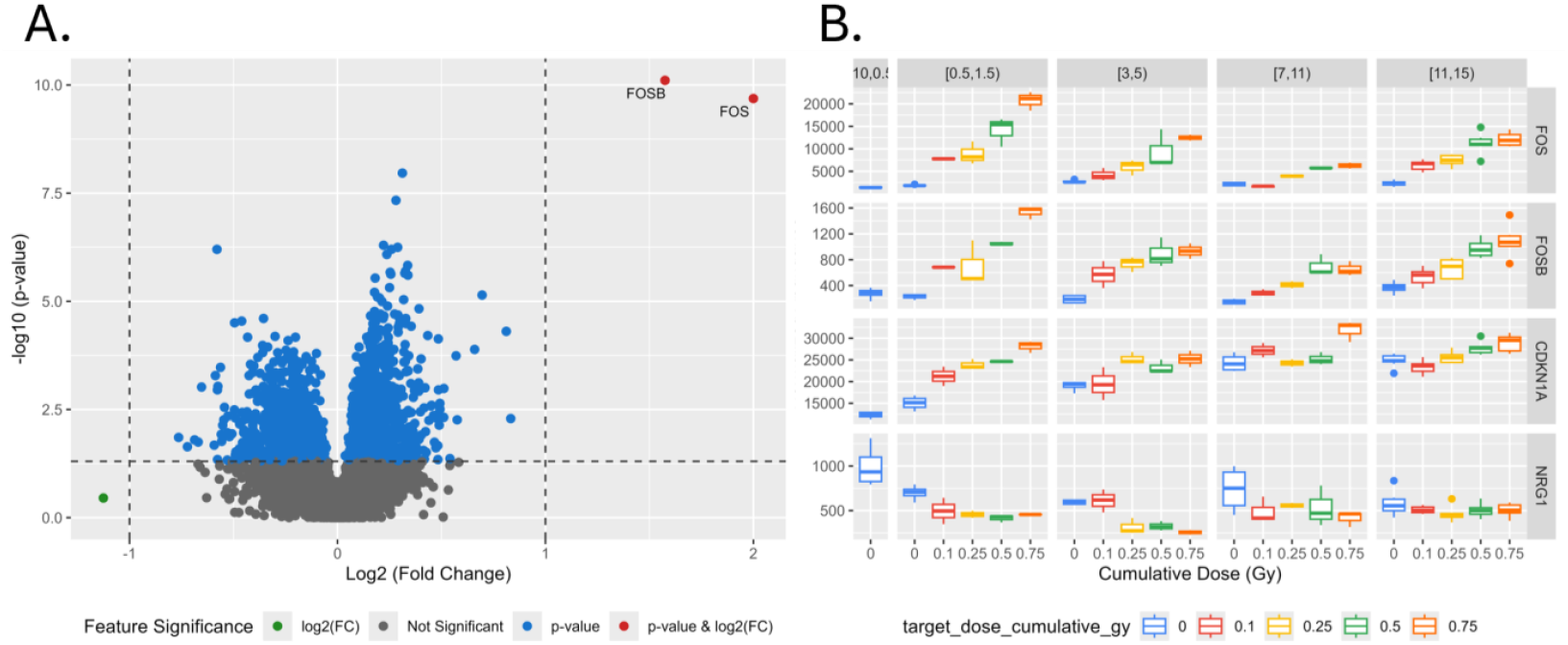
Differentially expressed genes in neutron irradiated EpiDermFT samples. **(A)** Volcano plot of RNA-seq-identified genes. Each gene is represented by a dot. Genes with negative log_2_ fold-change showed higher expression in the sham samples, and genes with positive log_2_ fold-change showed higher expression in samples exposed to neutron radiation. **(B)** Boxplots showing the normalized abundances of *FOS, FOSB, CDKN1A*, and *NRG1* across sampling days (-1, 1, 4, 7, 11 and 14) and radiation dose (0.0, 0.10, 0.25, 0.50, and 0.75 Gy).

### Gene Ontology (GO) Term Enrichment Analysis

The significantly up- and down-regulated genes identified from DESeq2 were converted to GO terms and secondary enrichment analysis was performed with only a few GO terms being enriched in the collection of significantly up-regulated genes (Figure 2). The enriched GO terms in the up-regulated genes included GO:0031575 (mitotic G1/S DNA damage checkpoint signaling), GO:0044819 (mitotic G1/S transition checkpoint signaling), and GO:0097305 (response to alcohol) (Figure 2A). More GO terms were enriched in the down-regulated genes; 15 GO terms were enriched in total (Figure 2A). Four main groups of enriched GO terms emerged in the set of significantly down-regulated genes (Figure 2B). Enriched GO terms were associated with either nucleosome regulation (GO:0034728, GO:0006334, GO:0000786, GO:0030527, and GO:0065004), ossification (GO:0030282, GO:0031214, GO:0002063, GO:0001503, and GO:0045071), DNA replication (GO:00090329, GO:0006261, GO:0006260, and GO:0030174), and one GO term associated with viral genome replication (GO:0045071). The GO term with the highest enrichment in both the up and down-regulated genes was GO:0030174 (regulation of DNA-templated DNA replication initiation) with a fold enrichment of 22.66. The complete break-down of the GO term enrichment can be found in supplemental Table S3.

**Figure 2:**
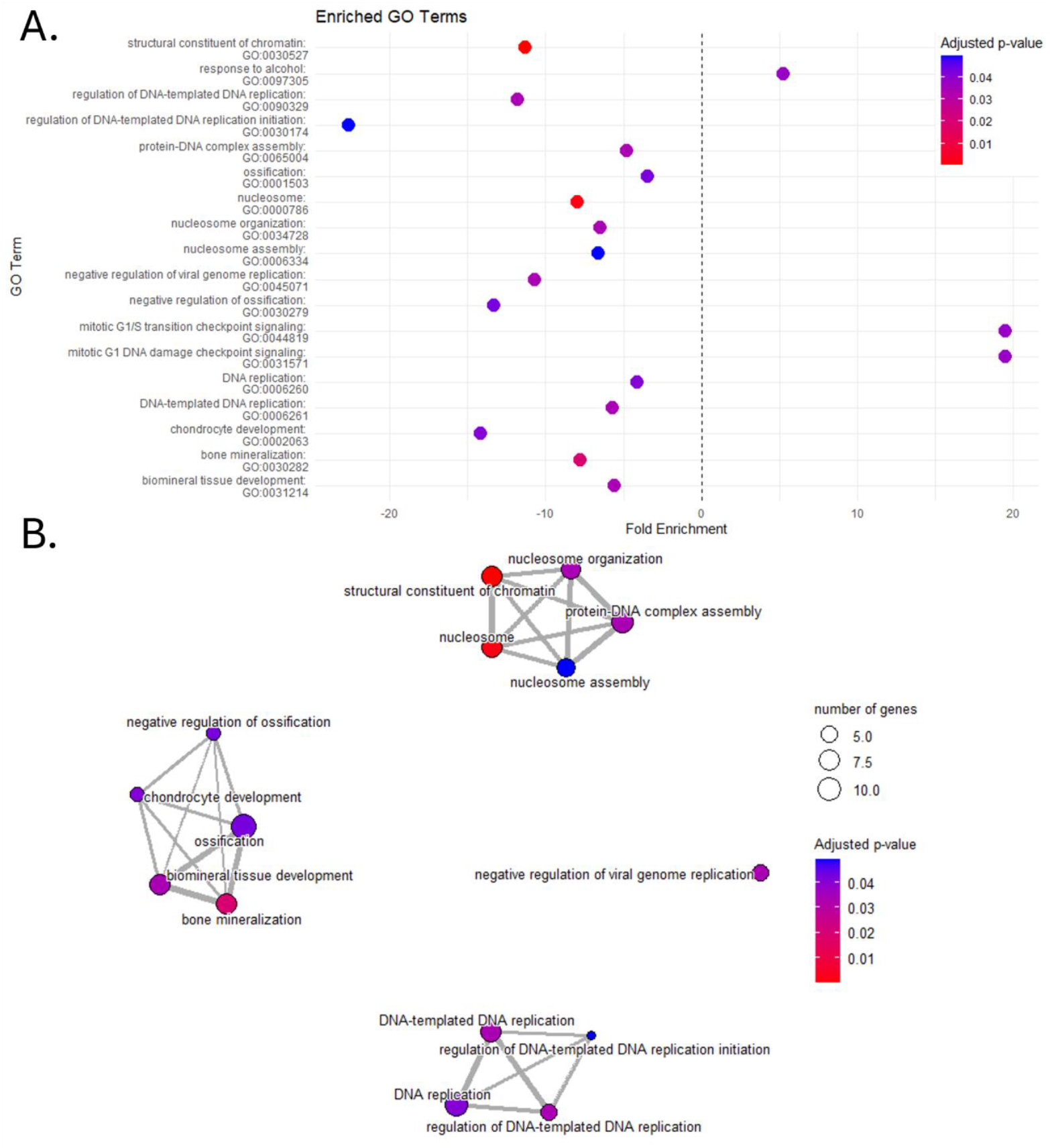
GO Term Enrichment plot. GO term enrichment was performed on significantly up- and down-regulated genes identified in the differential expression analysis **(A)**. Significantly enriched GO terms were determined after converting the gene ids to GO terms and comparing those to the normal background GO terms in the whole dataset. Significance was determined using the following parameters: p-value threshold of 0.05, q-value threshold of 0.05, and adjusted p-value less than 0.05 after false discovery rate multiple test correction. The GO terms from the down-regulated genes are plotted as an enrichment network map **(B)** showing the relationship between the number genes that share the same GO terms.

Of the166 down-regulated genes, 35 contributed to the 15 enriched GO terms (Figure 3A). Many of the genes did stay within the larger GO term cluster as shown in Figure 2. For example, eight histone protein genes were identified and were only associated with nucleosome assembly functions. However, there were some genes that had more diverse cross functionality and contributed to the GO term enrichment across the different clusters. *CDT1* and *CENPX* play a role in both the nucleosome organization and assembly, as well as DNA replication. Similarly, *ISG15* and *IFITM1* contribute to the enrichment of the negative regulation of viral genome replication as well as several ossification related GO terms. In the enriched GO terms from the up-regulated genes, 11 of the 127 genes were associated with the enriched GO terms discussed prior. Two genes, *CDKN1A* and *MDM2*, were attributed to all the enriched GO terms. Notably, *FOS* and *FOSB* contributed to the “response to alcohol” GO term, but *CDKN1A* contributed to the enrichment of all the enriched functions. *CDKN1A, FOS*, and *FOSB* are some of the genes that exhibited strong correlation with sample exposure, cumulative dose, and day post-exposure (Figure 1B).

**Figure 3:**
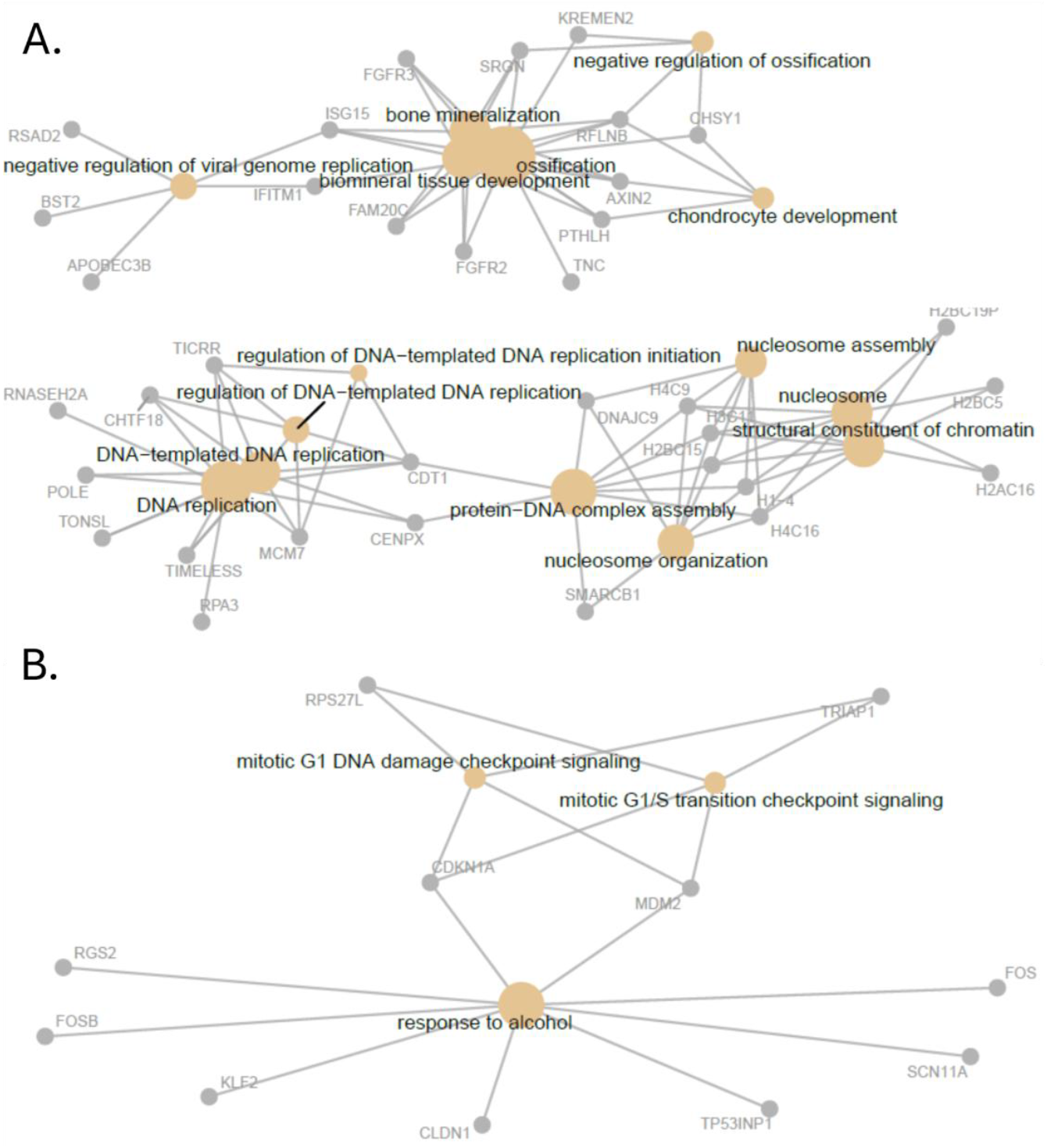
Cnet plot (gene-concept network) showing the significantly enriched GO terms (yellow dots) and their respective differentially expressed genes (gray dots) that contributed to the enrichment GO term. The edges connect each of the genes to their respective GO terms. **(A)** Includes the 15 enriched GO terms and 35 genes from the group of differentially down-regulated genes. **(B)** Includes the 3 enriched GO terms and the 18 genes from the differentially up-regulated genes.

### Predictive Modeling

Feature selection routines and manual review of the genes detected from RNA-seq resulted in 281 genes being selected to train machine learning models to predict whether samples had been exposed to neutron radiation, the cumulative radiation dose, and days post-exposure. For the comprehensive list of features and their use in the respective models, see supplemental Table S4. Figure 4 shows results for the top-performing models in each of these tasks, and performance metrics for each of the model types investigated are included in Table 3.

**Figure 4:**
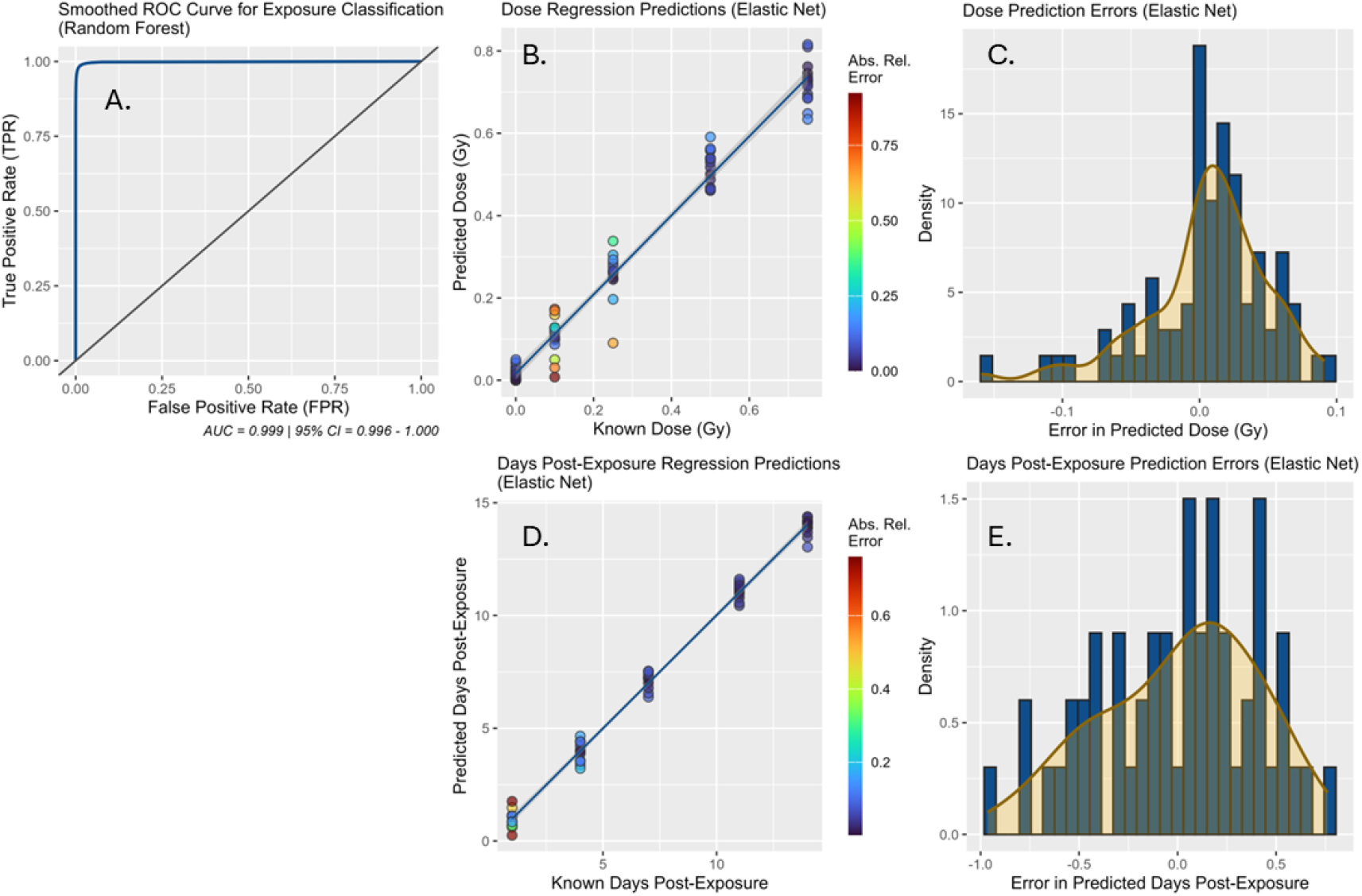
Results of the top-performing models (cross-validation resampled). **(A)** ROC curve for the Random Forest model predicting whether samples were exposed to neutron radiation. **(B)** Regression plot and **(C)** error histogram for the Elastic Net model predicting radiation dose. **(D)** Regression plot and **(E)** error histogram for the Elastic Net model predicting days post-exposure.

Classification models could consistently predict if samples had been exposed to neutron radiation with high accuracy. Random Forests was the top-performing model with an AUC of 0.9995 from repeated cross-validation results indicating near-perfect performance (Figure 4A), and other types of classification models had comparable results. These results suggest that even at doses of 0.1 Gy, there were marked differences in gene expression in our feature panel that were reliably diagnostic of exposure through 14-days post-exposure.

Next, we trained regression models to predict the radiation dose. All model types had similar success in this task with R_2_ values over 0.94 and the Elastic Net model achieving the highest R^2^ value of 0.9668 (Figure 4B and C). Although this model’s predictions generally had higher relative errors at lower doses, it maintained a low overall mean absolute error (MAE) of 0.0368 Gy. The other regularized linear regression model trained for this task, glmnet, was the second-best performer, outperforming the support vector machine (SVM) and neural network models—further indicating the linear relationships of these models were largely successful in capturing the dose effects present in our data.

Finally, days post-exposure was modeled in the samples exposed to neutron radiation using the same set of regression model algorithms. As with the dose predictions, the Elastic Net model performed the best, with an R^2^ value of 0.9921 and an MAE of 0.3482 days (Figure 4D and E). The other models performed similarly, all having MAE values under 1 day (Table 3).

## Discussion

In this study, we used RNA-seq to measure the gene expression response in an *in vitro* human skin model exposed to low dose neutron radiation over a two-week timeframe and used this transcriptomic data to develop machine learning models to predict exposure, dose, and days post-exposure. Our results show that these RNA signatures serve as reliable quantitative indicators of neutron radiation that enable both the classification of irradiated samples and regression-based estimation of the dose and time since exposure.

The high-dimensional gene expression data provided by RNA-seq is a powerful tool for biodosimetry efforts. Although previous biodosimetry studies have used RNA-seq to analyze transcriptomic responses to X-ray radiation,^25,26^ neutron biodosimetry has relied more on arrays and fixed gene panels.^27,28^ Unlike targeted approaches, which are limited to previously known biomarkers, RNA-seq offers an unbiased survey of the transcriptome and provides an expanded feature space for down-stream machine learning applications. Our machine learning pipeline implemented multiple feature selection strategies to identify the most informative gene panels prior to predictive modeling. One benefit of the large feature panels in the present study is that they enabled accurate dose prediction even when the time point of irradiation (i.e., days post-exposure) was not known.^29^ Gene expression changes dynamically after radiation exposure, which complicates accurate dose prediction, as individual genes may show similar upregulation at different time points depending on the dose of radiation. By utilizing a larger panel of genes, each with their own response kinetics, the regression models were able to accurately determine the dose of radiation.

Regression model predictions for days post-exposure were accurate, with the Elastic Net model achieving an R^2^ value of 0.9921. While this performance may reflect genuine transcriptional differences across stages of the radiation response, it is also likely that the models captured variation related to the degradation of tissue samples over time. This tissue degradation may have been associated with changes in gene expression that coincided with days post-exposure. Future data analyses are required to determine the degree to which gene kinetics in the neutron radiation response may be used to predict time post-exposure.

### Differential Expression Analysis

After exposing samples to neutron radiation and performing the RNA extractions, sequencing, and bioinformatics processing, we performed differential expression analysis on the data. We detected approximately 21,535 genes in total and of those 832 genes showed statistically significant changes in regulation; however, with less permissive selections there were only 320 genes that were statistically significant and had an absolute log_2_ fold change greater than 0.25.

We then performed the differential expression analysis for each dose and day post-exposure compared to the controls, from which we observed distinct trends when looking at the number of significantly expressed genes in relation to dose and days post-exposure. Regarding dose, there appeared to be a dose dependent jump where the lower Gy (0.1 Gy: n = 49 and 0.25 Gy: n = 33) yield fewer differentially expressed genes compared to the higher doses (0.5 Gy: n = 100 and 0.75 Gy: n = 93). Interestingly many of the genes that show strong predictive power only appear to only be differentially expressed after 0.1 Gy. For example, *MDM2, TP53INP1*, and *NRG1* show signs of significant differential abundance only after 0.25 Gy (see supplemental Table S2). In addition, the differentially expressed genes at 0.1 Gy were not well conserved across the other doses. Only *EMILIN3, FOS, FOSB*, and *PRKX* were expressed at 0.1 Gy and at least one other dose level, whereas 26 of the 33 genes in the 0.25 Gy dose were conserved in at least one of the other doses. Eleven (11) of the 33 genes found in 0.25 Gy were conserved in both 0.5 and 0.75. These include *KLF6, CCNG1, FOSB, FOS, MDM2, GNAL, NRG1, TP53INP1, TRIAP1, PRKX*, and *RPS27L*.

Differentially expressed genes showed little uniformity across the varying doses, suggesting a stronger temporal association between radiation exposure and gene expression. Our results show that there is a substantial increase in the number of differentially expressed genes on day 4 and day 11 post neutron exposure but then reduces to as low as 16 genes after 14 days. We detected a total of 554 unique genes that had differential expression when considering each timepoint, but only 29 were conserved across more than one timepoint. Similar to cumulative dose, *FOS* and *FOSB* were the only genes conserved across all the time points. Our results suggest there is a lag time between exposure and when the limited conserved changes in expression are observed. Of the 29 conserved genes, 15 were observed on both day 4 and day 11 post-exposure. Interestingly, several of these 15 genes showed a transition from positive to negative log_2_ fold change. For example *MKI67*, a gene associated with sun damaged skin tissue,^30^ was down-regulated at day 4 but was up-regulated at day 11. This is possibly due to the role *MK167* plays in various cancer types, increasing immune cell infiltration into cancer cells as well as controlling the cell cycle.^30,31^ *MK167* downregulation is associated with cell cycle arrest in conjunction with *p53* and indicates cell cycle arrest; therefore, seeing downregulation earlier post-exposure is not unsurprising. The reversal *MK167* expression, becoming significantly up-regulated by day 11 suggests the tissue is attempting to return to normal cellular function.

Taken collectively, these temporal results corroborate existing studies, like those performed by Qutob and colleagues in 2024 in which they observed a significant increase in the number of differentially expressed genes at 6 and 48 h post-UV exposure at both 20 and 40 mJ/cm^2^.^32^ Qutob et al. also showed that after 72 and 96 h post exposure at 20 and 40 mJ/cm^2^ respectively, there was a substantial decrease in the number of differentially expressed genes. Even though our results were generated using a different tissue model and using a different ionizing radiation source, we saw similar temporal fluctuations in gene expression. We detected a substantial jump in the number of differentially expressed genes at day 4 (n = 255) and then a large decrease in expression at day 14 (n = 16) (see Table 2). This could be a similar phenomenon in which the cells activate numerous genes to repair the damage and then return to a basal level. However, we cannot definitively say this was the case, as by day 14 the tissue samples were beginning to degrade, and our confidence is low that the lack of differentially expressed genes was the result of a return to baseline metabolism. It is likely that the cellular degradation caused further cellular damage and the expected genes were activated in the control samples regardless of radiation exposure. This may be why in Figure 1B, the initial down-regulation of the growth factor *NRG1* dissipates after the first few days.^33^ We postulate that the exposed samples had a decrease in *NRG1* expression to inhibit cell growth, allowing time for DNA repair. Conversely, the sham control samples were able to continue proliferating during early timepoints but eventually arrested at later timepoints, likely due to the accumulation of age-related damage. If the exposed samples were returning to normal, we would expect to see the up-regulation of genes, like *NRG1*, to the levels observed in the control samples at day -1 and day 1. Instead, *NRG1* showed a decreasing average read count over time in the sham samples and no change in the exposed samples. This observation suggests that regardless of exposure status, cells were not returning to normal cell growth.

### GO Term Enrichment

GO term enrichment analysis of the up- and down-regulated genes provides a global overview of changes in cellular activity compared to individual genes. We noticed three major changes in the enriched GO terms for the up-regulated genes: “response to alcohol”, “mitotic G1/S DNA damage checkpoint signaling”, and “mitotic G1/S transition checkpoint signaling” (GO:0097305, GO:0031575, and GO:0044819 respectively; see Figure 2). Many of the genes contributing to “response to alcohol” included *MDM2, CDKN1A, TP53INP1, FOS*, and *FOSB*, all of which were highly predictive features for our biodosimetry models. Initially, the enrichment of the “response to alcohol” seems unusual. However, both alcohol exposure and radiation exposure promotes the formation of reactive oxygen species.^34,35^ Therefore, it is likely that the “response to alcohol” enrichment is simply due to the increase in reactive oxygen species caused by the radiation exposure. The other two GO terms, “mitotic G1/S DNA damage checkpoint signaling” and “mitotic G1/S transition checkpoint signaling” are not unusual. *MDM2* and *CDKN1A* often work in concert to control cell proliferation. As cells are exposed to stress, like radiation, there are higher levels of *CDKN1A* and *MDM2* to control cell arrest to allow time for cells to repair any damage to the DNA.^36,37^

An unexpected result was the enrichment of ossification GO terms, considering the model was epidermal. Bone loss has been reported in response to radiation exposure in multiple studies.^38–40^ Though skin and bone tissue seem disparate, the down-regulated genes, *AXIN2, PTHLH* and *RFLNB*, also possess functions in the both tissue types. *AXIN2* acts as a negative regulator for Wnt pathways to prevent overexpression and subsequent tumor formation. *PTHLH* is down-regulated post radiation exposure leading to hair loss but is also critical in cartilage and bone formation.^41,42^ Furthermore, *PTHLH* produces PTHrH, a protein hormone that controls cell proliferation and differentiation across multiple different tissue types including bone, cartilaginous, and skin.^43^ *RFLNB*, or refilin B, is a protein associated with actin bundle organization as well as bone mineralization regulation during development. In this study it is likely that the exposure to radiation influenced cytoskeletal reorganization, as there has been prior documentation of cell morphology changes at low levels of ionizing radiation exposure.^41,44,45^ More investigation is necessary to better understand what overlap exists between the ossification GO terms and skin tissue, but this is likely due to an overlap between hair growth related pathways and bone formation during early cellular differentiation.

The decrease in GO terms related to nucleosomes and DNA replication in irradiated skin tissue is a direct consequence of the cell cycle arrest and DNA damage response mechanisms activated by ionizing radiation. Radiation induces widespread DNA strand breaks, forcing cells, particularly the rapidly dividing stem and progenitor cells in the epidermis to immediately halt their proliferation. The cell cycle arrests to allow for DNA repair mechanisms to engage, or if damage is too extensive, to undergo apoptosis. Lastly, the decrease in expression of *RSAD2, BST2, APOBEC3B* and the subsequent enrichment of the “negative regulation of viral genome replication” is a well-documented response to radiation and not surprising. It has been well characterized that host innate immune responses suffer after radiation exposure, making exposed individuals more susceptible to.^46,47^ The radiation doses in our study were relatively low; however, neutron radiation is highly damaging and so the low Gy exposure is likely to still cause severe damage to the tissue.

## Conclusion

This study demonstrates that RNA sequencing of a 3D *in vitro* human skin model enables sensitive characterization of low-dose neutron radiation exposure over an extended post-exposure window. Across doses up to 0.75 Gy and sampling through 14 days post-exposure, we identified reproducible transcriptomic signatures associated with DNA damage signaling, cell cycle arrest, chromatin regulation, and cellular stress responses. The core genes identified included *FOS, FOSB, CDKN1A, MDM2*, and *GADD45A* consistently reflected the exposure characteristics, and the larger set of genes align with previously published coordinated biological responses to radiation exposure.

Machine learning models trained on these transcriptomic features accurately classified exposed versus unexposed samples and reliably estimated the radiation dose, even at low cumulative exposure levels. These results support the utility of skin-derived RNA signatures for biodosimetry applications where exposure history is unknown and samples may be collected non-invasively and days after an incident. Although regression models predicting days post-exposure exhibited very high accuracy, these predictions were likely confounded by progressive degradation of the *in vitro* tissue model at later time points. This limitation highlights the need to separate radiation-specific temporal biological responses from tissue aging effects in future studies.

Overall, this work establishes proof of principle that transcriptomic signatures from human skin tissue can support quantitative, non-invasive assessment of neutron radiation exposure, providing a foundation for future translation to real-world sample types and complex exposure scenarios.

## Acknowledgements

The authors would like to thank Dr. Sungyeon Jang, Dr. Lydia Contreras, and the University of Texas Genomic Sequencing and Analysis Facility for their help with RNA sample submission and RNA sequencing data acquisition. This research was supported by the Intelligence Advanced Research Projects Activity (IARPA) Targeted Evaluation of Ionizing Radiation Exposure (TEI-REX) program through the Army Research Office (ARO) contract No. W911NF22C0051. The views and conclusions contained should not be interpreted as necessarily representing the official policies, either expressed or implied, of ODNI, IARPA, ARO, or the U.S. Government. The U.S. Government is authorized to reproduce and distribute reprints for governmental purposes notwithstanding any copyright annotation therein. This paper was typeset with the bioRxiv word template by @Chrelli: https://github.com/chrelli/bioRxiv-word-template.

## Author contributions

MWG - conceptualization, data curation, supervision, methodology, software, writing – review & editing, formal analysis; DSL - conceptualization, methodology, investigation, sample preparation and data collection, writing – original draft, writing – review & editing, visualization.; JCG - data curation, formal analysis, writing – original draft, writing – review & editing, visualization; JCW - writing – original draft, writing – review & editing, software, validation, data curation, formal analysis, visualization; MNP - methodology, investigation, sample preparation and data collection, formal analysis; GG - resources, methodology; EAS - writing – review & editing, methodology, investigation, sample preparation and data collection; HCT - resources, methodology.

PJC: software, validation, formal analysis, visualization, data curation. CG: investigation and sample collection; VMJ - sample preparation and data collection; CAV - sample preparation and data collection; BET - sample preparation and data collection; FCH - conceptualization, project administration, writing – review & editing.

## Materials and Methods

### Maintenance of Human *In Vitro* Skin Tissue Culture

MatTek EpiDerm Full Thickness (FT) skin model samples were shipped directly from MatTek in sealed 6-well plates. Upon receipt samples were stored in their unopened 6-well plates at 4 °C for no longer than 24 hours. Day 1 began with the aseptic addition of EFT-400 maintenance medium (EFT-400-MM) that had been pre-warmed to 37 °C to the total number of wells required for the study. Sterile forceps were used to transfer EpiDermFT-400 tissues in their inserts to 6-well plates pre-filled with EFT-400 media. A total of 84 samples were used for the study; these samples were spread across fourteen (14) 6-well plates. The tissues were equilibrated in their new 6-well plates at 37 °C and 5% CO2 overnight.

On Day 2 and every day of the study thereafter, the spent media in each EpiDermFT sample well was aspirated and replaced with fresh, pre-warmed (i.e., 37 °C) EFT-400-MM. Any media on top of the tissues was carefully removed. The tissues were incubated at 37 °C and 5% CO2. Following the media change on Day 2, samples were irradiated. Media was changed daily following irradiation.

### Neutron Radiation Exposure

Neutron irradiation of all sample types was performed using the Columbia IND Neutron Facility (CINF) located at the Radiological Research Accelerator Facility (RARAF). The neutron field was produced by a mixed beam, composed of 5 MeV atomic and molecular ions of hydrogen and deuterium that was used to bombard a thick beryllium (Be) target. One or two sample plates were placed on a shelf 190 mm from the target center and at an angle of 60 degrees from the ion beam direction. To provide uniform irradiation, sample plates were flipped front-to-back halfway through the irradiation.

The dosimetry for the irradiations was performed on the morning of the experiment using a custom tissue-equivalent (TE) gas ionization chamber, placed on the sample holder wheel. This chamber measured the total dose in the mixed neutron and γ-ray field. To evaluate the ratio of neutron and γ doses, gamma-ray dosimetry was performed separately by replacing the ionization chamber with a compensated Geiger-Mueller dosimeter, which has a very low neutron response. Overall dose uncertainty is ±5%, primarily due to uncertainties in kerma (kinetic energy released per unit mass) and mass stopping power corrections used to convert from the ionization chamber measurement to dose. Typically, the field consisted of approximately 20% (by dose) photons and 80% neutrons. Details regarding the neutron irradiation setup are in Supplemental Materials (Table S5). The actual doses delivered to each plate are shown in Table S6.

### EpiDermFT Sample Collection

To collect the samples, each EpiDermFT tissue sample was removed from its well and placed onto a sterile work surface (e.g., sterile petri dish). The bottom cap of the insert was aseptically removed to release the EpiDermFT tissue sample from the plastic housing, making sure the insert membrane is no longer attached to the tissue. A No. 10 scalpel was then used to section the EpiDermFT tissue sample into quarters. Each quarter was transferred into a microcentrifuge tube containing 300 µL of RNAlater (Invitrogen). Samples were collected prior to radiation exposure (day -1) and on days 1, 4, 7, 11, and 14. The EpiDermFT samples were stored at -80 °C until extraction. Sample metadata can be found in the Supplemental Material (Table S7).

### EpiDermFT Sample Extraction

Samples stored in RNAlater were pre-processed by removing samples from the RNA preservative, adding 300 µL 1X PBS in a Lyse&Spin basket (Qiagen), and spinning for 1 minute at 20,000 × g to remove residual RNAlater. Washed quarters were placed into ice chilled ZR Bashing-Bead 2.0 mm lysis tubes (Zymo) with 600 µL Qiagen Buffer RLT Plus with added β-mercaptoethanol. Samples were lysed at room temperature for 3 minutes at 3,000 rpm using a Cell Disruptor Genie (Scientific Industries). RNA was extracted from lysed samples using the Qiagen RNeasy Plus Mini Kit per manufacturer instructions. RNA concentration was determined using the Invitrogen Qubit RNA Broad Range kit. Details regarding the various extraction methods tested can be found in the Supplemental Methods (Table S8).

### RNA Sequencing of EpiDermFT

Sequencing was performed on RNA samples extracted from EpiDermFT tissue sections by the University of Texas Genomic Sequencing and Analysis Facility (GSAF). After RNA extraction and Qubit analysis, a fraction of the eluted RNA was diluted in nuclease-free water to meet GSAF sequencing input requirements of at least 250 ng total RNA in a volume of at least 25 µL. If the sample was below the 250-ng threshold the entire sample volume (25 μL) was provided. A total of 84 samples were submitted for sequencing, with 35 samples at a concentration above 25 ng/µL of RNA, 45 samples at a concentration below this threshold, and four samples that had approximately 0 ng of RNA. Sequencing samples were prepared in 96-well plates and stored at - 80 °C prior to sequencing library preparation. Low volume samples were prepared in separate 96-well plates, according to the University of Texas GSAF guidelines.

RNA sequencing libraries were assembled using the NEBNext Ultra II RNA Library Prep Kit for Illumina. Following library preparation, library quality was assessed on an Agilent 2100 Bioanalyzer using a Bioanalyzer DNA 1000 chip. Sequencing was performed on an Illumina NovaSeq with minimum of 20,000,000 reads and a target of 25,000,000 reads.

### Bioinformatic Analysis of RNA-seq Data

Raw paired end RNA sequences were trimmed using FaQCs^48^ v2.10 using a minimum quality cutoff of 20, and a minimum length of 50 BP for each read. Trimmed reads are quantified and mapped to host genomes using Kallisto^49^ v0.50.1. The host genomes used was HG38 for EpiDermFT tissue cultures. Gene names are converted to Ensembl gene names. Differential gene expression analysis of the trimmed reads was performed using DESeq2^50^ v1.44.0. Batch correction was performed using ComBat-Seq^51^ v3.52.0. Reads were normalized using median of ratios method and updated sequence count data were generated with statistical analyses performed on all extant mapped genes. Multiple test corrections were applied to p-values using false discovery rate (FDR).

### RNA Sequencing Data Analysis and Predictive Modeling

After raw sequences were processed, differential gene expression was calculated using DESeq2. Ensembl gene IDs were converted to the HUGO Gene Nomenclature Committee (HGNC) term and reported throughout. Sham samples were compared to all exposed samples and were treated as the single experimental group. Mean read count, log_2_ fold change, lfcSE (standard error of the log_2_ fold change), Wald statistic, p-value, and adjusted p-value using false discovery rate was calculated for every gene detected. The same parameters were applied when performing the differential expression analysis for each specific dose to determine how gene expression changes in relation to cumulative dose.

Differentially abundant genes with an absolute log_2_ fold-change greater than 0.25 and an adjusted p-value less than 0.05 were grouped into two groups, up- and down-regulated genes. These genes’ normalized read counts were pulled and the ENSEMBL gene id was converted to ENTREZ IDs. ClusterProfiler^52^ v4.12.6 was then used to perform the GO term enrichment analysis identifying both parent and child GO terms that were enriched in the up- and down-regulated genes. Multiple test corrections were applied to p-values using FDR and cutoff for p-values and q-values were both less than 0.05.

Genes used for model development had an adjusted p-value less than 0.05 and a mean read count greater than 500 reads. The genes that had an absolute log_2_ fold change less than 0.25 were removed. Boxplots were generated showing the distribution of read counts for each remaining gene across sampling time and dose. If there were insufficient candidates, the log_2_ fold change threshold of 0.25 was removed and anything greater than 0 was considered.

Two machine learning methods, Lasso regression and the Boruta^53^ algorithm, were used to select features (genes) for predictive modeling. These methods were applied in a model-specific manner, identifying three separate panels of features each to be used for modeling a different outcome variable (exposure, dose, and days). The panels of features used for modeling exposure and dose, in addition, included all genes identified from differential expression analysis. Boxplots were generated for all features selected by this approach and manually reviewed for final inclusion in its respective feature panel.

Based on principal component analysis (PCA), two samples were identified as outliers and excluded from further analysis. After removing these samples, some features were still observed to have significant outliers or biologically implausible distributions and were considered anomalies. Supervised anomaly detection was used to identify and remove these features based on their distributions across samples. The remaining feature data were mean-centered and scaled to unit variance prior to feature selection and modeling.

Model development and evaluation was conducted using RStudio Server with R version 4.4.2 and utilized functions from the caret (Classification and Regression Training) package^54^ version 7.0-1 for model training. Separate models were developed for predicting each outcome: classification models for predicting if samples had been exposed, and regression models for predicting the radiation dose and days post-exposure. Models for predicting exposure and dose were developed using all samples, whereas models for predicting days post-exposure were developed using only exposed samples.

Models of various algorithms for classification and regression (e.g., logistic regression, boosted logistic regression (LogitBoost), Random Forests,^55^ Support Vector Machine with linear or radial kernels,^56^ Generalized Linear Models with Elastic Net Regularization (glmnet)^57^, ElasticNet (enet), and Bayesian regularization for feed-forward neural networks (brnn) were fit and evaluated for each of these tasks with 10 repeats of 10-fold cross-validation in which 80% of the samples were used for training and 20% were used for validation in each repeated fold. All modeling results presented are from the repeated cross-validation performance.

## Supplemental Material

### Supplemental File Attachments

**Table S1: Differential gene expression analysis results**.

The complete results from the DESeq2 differential expression analysis of sham versus exposed samples.

**Table S2: Dose and Day specific DESeq2 differential expression analysis**.

Results containing trimmed DESeq2 results with only genes with adjusted p-values less than 0.05 and an absolute log2 fold-change greater than 0.25 at each specific dose and day post exposure. Included in the trimmed table is the protein name, ENSEMBL gene id, log_2_fold-change and dose.

**Table S3: Gene Ontology (GO) term enrichment analysis**.

Results from the GO enrichment analysis for both the upregulated and downregulated statistically significant differentially expressed genes.

**Table S4: Modeling features**.

Genes used as features for each predictive model.

**Table S7: Sample metadata**.

Metadata file with sample information used for data analysis.

## Supplemental Tables

### Detailed Neutron Irradiation Conditions and Radiation Doses Samples Received

**Table S5:** Detailed neutron irradiation conditions.

| Parameter | Value |
| --- | --- |
| Source Type | Broad spectrum neutrons |
| Generation Mechanism | Accelerator (10 μA of a 5 MeV mixed beam of protons and deuterons impinging on a thick beryllium target). The irradiation setup and dosimetry are described in Xu et al., 2015. |
| Dose Rate | 1.1 Gy/h |
| Source-to-Target Center Distance | 190 cm |
| Detector Type | Tissue equivalent proportional counter (TEPC; total dose) as described in Rossi et al., 1960. Compensated Geiger Müller counter (GM; photon dose) |
| Dosimetry Setup | Both GM and TEPC are irradiated at the same position as the mice and used daily to calibrate a monitor TEPC placed at a fixed location. Beam delivery is controlled by the monitor |
| Calibration Plan | Dosimetry is calibrated to a <sup>226</sup>Ra seed before each experiment. Currently each count on the TEPC electronics chain corresponds to 11 nGy of absorbed dose to tissue |

**Table S6:** Neutron irradiation doses received by EpiDermFT samples.

| Plate | Collection Days | Neutron Dose (Gy) | Gamma Dose (Gy) | Total Dose (Gy) | Dose Rate (Gy/h) | Duration (s) |
| --- | --- | --- | --- | --- | --- | --- |
| 1 | D7 / D11 | 0.60 | 0.15 | 0.75 | 1.1 | 2,454 |
| 2 | D14 | 0.60 | 0.15 | 0.75 | 1.1 | 2,454 |
| 3 | D7 / D11 | 0.40 | 0.10 | 0.50 | 1.1 | 1,636 |
| 4 | D14 | 0.40 | 0.10 | 0.50 | 1.1 | 1,636 |
| 5 | D7 / D11 | 0.20 | 0.05 | 0.25 | 1.1 | 818 |
| 6 | D14 | 0.20 | 0.05 | 0.25 | 1.1 | 818 |
| 7 | D7 / D11 | 0.08 | 0.02 | 0.10 | 1.1 | 327 |
| 8 | D14 | 0.08 | 0.02 | 0.10 | 1.1 | 327 |
| 9 | D1 / D4 | 0.60 | 0.15 | 0.75 | 1.1 | 2,454 |
| 10 | D1 / D4 | 0.40 | 0.10 | 0.50 | 1.1 | 1,636 |
| 11 | D1 / D4 | 0.20 | 0.05 | 0.25 | 1.1 | 818 |
| 12 | D1 / D4 | 0.08 | 0.02 | 0.10 | 1.1 | 327 |

## Supplemental Methods

### EpiDermFT Sample Extraction Workflow Development

To determine the best approach for extraction, 12 EpiDermFT tissues were maintained and collected as described in the main Methods, with all samples collected on day -1. This resulted in 48 total sections. Upon collection, samples were stored in either 300 or 600 µL Zymo DNA/RNA shield or 600 µL Invitrogen RNAlater. Collected samples sat for approximately one hour at room temperature and then stored at -80 °C overnight prior to extraction. Samples stored in RNAlater underwent a PBS wash step, adding 300 µL 1X PBS in a Lyse&Spin basket (Qiagen), and spinning for 1 minute at 20,000 × *g*. Various extraction methods were tested including different storage conditions, lysis buffers, and lysis conditions (Table S1). Relevant equipment included a thermoshaker, an Omni Bead Ruptor (Omni International) with liquid nitrogen, and a Cell Disruptor Genie (Scientific Industries). If appropriate β-mercaptoethanol (βMe) was added as indicated. ZR BashingBead 2.0 mm lysis tubes (Zymo) were used for all non-heat shake lysis conditions. RNA concentrations were determined using the Qubit RNA Broad Range kit (Invitrogen) and RNA integrity number (RIN) by the Agilent 2100 Bioanalyzer using the RNA 6000 Pico kit (Agilent). Testing is described in the Supplementary Results section below. The best performing method was then used for all study samples as described in the main text.

**Table S8:**
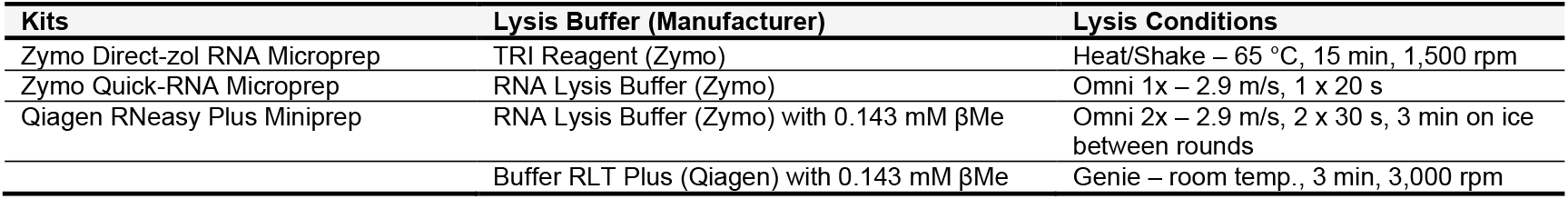
Details of tested RNA extraction conditions.

| Kits | Lysis Buffer (Manufacturer) | Lysis Conditions |
| --- | --- | --- |
| Zymo Direct-zol RNA Microprep | TRI Reagent (Zymo) | Heat/Shake – 65 °C, 15 min, 1,500 rpm |
| Zymo Quick-RNA Microprep | RNA Lysis Buffer (Zymo) | Omni 1x – 2.9 m/s, 1 x 20 s |
| Qiagen RNeasy Plus Miniprep | RNA Lysis Buffer (Zymo) with 0.143 mM βMe | Omni 2x – 2.9 m/s, 2 x 30 s, 3 min on ice between rounds |
| | Buffer RLT Plus (Qiagen) with 0.143 mM βMe | Genie – room temp., 3 min, 3,000 rpm |

## Supplemental Results

### EpiDermFT RNA Extraction Development

A small batch shipment of 12 EpiDermFT tissues were maintained, and quarter sections were collected after ∼24 hours post-receipt, mimicking the Day -1 samples. Following collection, multiple extraction kits and lysis methods were tested (Table S8, Figure S1A). Each condition used two quartered sections from different tissues for replicate testing. Samples in RNA*later* underwent a pre-processing PBS wash step using Lyse&Spin baskets prior to lysis. All samples were then evaluated for overall RNA yields and RNA quality (RNA integrity number, RIN) following extraction and elution.

Extracted samples were assessed against metrics needed for RNA-sequencing including a minimum of 500 ng total RNA yield and a RIN value of at least 7.0. This would provide high quality RNA for RNA-seq with enough total RNA for both RNA-seq and future RNA analysis methods if desired. It was noted that homogenization by Genie Disruptor (Scientific Industries) consistently gave higher yields across kits but often had remaining tissue material visible post-lysis. In contrast, samples processed by the Omni Bead Ruptor (Omni Inc) were fully homogenized with no visible tissue remaining but typically resulted in lower yields.

Results indicated two top performing methods, both using the Qiagen RNeasy kit (Figure S1B, boxed result). Both Qiagen-based methods had an average recovery above the minimum target yield and quality RNA. Other methods showed either low RNA recoveries or poor RNA quality, insufficient for the target sequencing goals. Qiagen RNeasy paired with the Genie Disruptor had reduced variability compared to the Omni Bead Ruptor approach, and this method was used for all study sample extractions.

**Figure S1:**
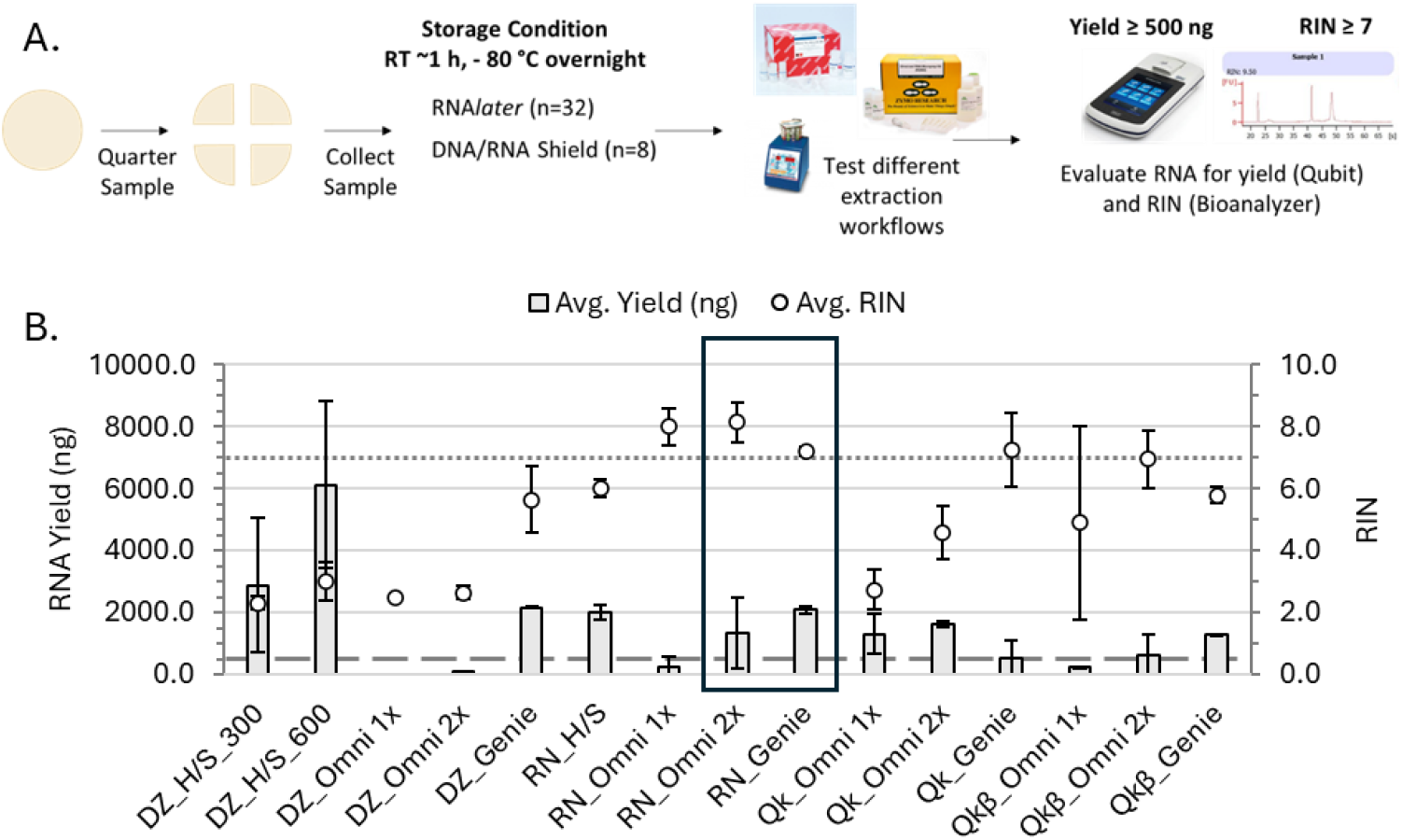
Extraction development results of EpiDermFT sections. **(A)** Workflow of RNA extraction testing. Quarter section samples were collected and RNA extracted using various storage, lysis, and kits to determine the best extraction workflow, using RNA yield and quality metrics **(B)** Results of EpiDermFT extractions using different workflows. DNA/RNA Shield samples are indicated with either 300 or 600; all other samples were stored in RNA*later*. The boxed data indicates the best performing conditions. The gray dashed line indicates the target RNA yield minimum of 500 ng and the gray dotted line indicates the target RNA quality (RIN) minimum of 7.0. Plots are the average ± SD (n=2). Abbreviations for kits and lysis conditions: Direct-zol RNA Microprep kit lysed in TRI Reagent, DZ; Quick-RNA Microprep kit lysed in RNA Lysis Buffer, Qk; Quick-RNA Microprep kit lysed in RNA Lysis Buffer with βMe, Qkβ; RNeasy Plus Miniprep kit lysis in RLT Buffer with βMe (RN); Heat/Shake, H/S. IDs are structured as: Kit/Lysis Buffer_Lysis Condition_Storage Solution (DNA/RNA Shield only). See also Table S8.

